# Selectivity of Lateral Epidural Spinal Cord Stimulation with Varying Electrode Contact Diameters and Stimulation Configurations

**DOI:** 10.64898/2026.06.14.732127

**Authors:** GJ Ansah, M Del Brocco, S Bhowmick, M Duran, C Gopinath, MK Jantz, SF Lempka, LE Fisher

## Abstract

***Objective*.** Our prior studies have demonstrated that lateral spinal cord stimulation can evoke somatosensory percepts in the missing foot in individuals with a lower-limb amputation. However, subjects reported concurrent sensations in their residual limb. In this study, we evaluate the hypothesis that using high-density paddle electrodes with smaller contact sizes, and multipolar stimulation configurations could evoke more focal sensations in the foot over a wide range of stimulation amplitudes. ***Approach.*** We used a combination of electrophysiology and computational modelling methods to investigate the selective activation of distal nerve branches in response to lateral spinal cord stimulation in cats. In six acute feline experiments, we performed an L3-S1 laminectomy and placed custom 32-electrode paddles laterally over the dura of the spinal cord. We recorded antidromic action potentials in the distal branches of the sciatic and femoral nerve trunks in response to stimulation using three contact diameters (150, 500 and 1000 µm) and two stimulation configurations – monopolar and bipolar stimulation. We replicated the neural recruitment patterns from those experiments in a computational model of the feline lumbar spinal cord. We then used the model to examine neural recruitment with 1.8 mm and 2.5 mm contacts, as well as a tripolar guarded-cathode configuration. ***Main results.*** In the electrophysiology experiments, the 500 µm-diameter electrodes achieved the most selective nerve activation (68%) compared to 62% for both 150 and 1000 µm-diameter electrodes. The minimum amplitudes for recruiting nerve branches (i.e., threshold) as well as the dynamic ranges were largely similar for the different contact diameters (median: 35 µA) and stimulation configurations (30 µA for bipolar stimulation; 35 µA for monopolar stimulation). The computational model reproduced the finding that selectivity did not differ significantly among the three contact sizes tested in cat experiments, though it revealed that increasing contact diameter above 1000 µm raised the minimum amplitude required for selective activation and reduced spinal root selectivity. Across both approaches, we consistently recruited large-diameter afferents that are critical for somatosensory applications of spinal cord stimulation. ***Significance.*** Our results indicate that, relative to clinical electrodes, reducing the contact diameter of stimulation electrodes can evoke focal sensations, but further reductions below 1000 µm may fail to improve selectivity. This study highlights potential constraints with achieving focal selectivity that are not dependent on the design of the electrodes.

## Introduction

The prevalence of lower-limb amputation is predicted to reach over 2.3 million people in the United States within the next decade^1^. Most people living with a lower-limb amputation rely on prosthetic devices for mobility. However, prosthesis users report several unmet needs associated with current state-of-the-art lower-limb prostheses, including an increased risk of falls, impaired mobility and balance, and pain in the residual limb and phantom limb^2^. Studies have shown that all these unmet needs may be associated with the loss of somatosensory feedback that occurs with limb amputation^3^. Therefore, restoring naturalistic tactile feedback in the missing limb has emerged as a promising approach to address the needs of people with lower-limb amputation^3–7^.

In a recent study conducted by our research group, we demonstrated in three individuals with transtibial amputation that lateral spinal cord stimulation (LSCS) can evoke sensations in the missing limb. In that study, LSCS was delivered via commercially available spinal cord stimulation electrodes, which are typically employed for alleviating chronic pain. Stimulation intensity was modulated in real time based on pressure signals recorded from a wireless pressure-sensing shoe insole so that the evoked sensations appeared to emanate from the prosthetic foot^8^. Our results showed significant benefits, including improved functional use of lower-limb prostheses, enhanced gait and postural stability, and a reduction in phantom limb pain. Despite these positive outcomes, we noted that the sensations evoked in the missing limb by LSCS were invariably accompanied by sensations in the residual limb^8^. These residual limb sensations can be distracting, since they co-occur with the sensations in the missing foot and do not provide additional meaningful information about the state of the prosthetic limb. Given our hypothesis that LSCS likely restores sensation by activating the densely packed, small-diameter dorsal rootlets (DR), which measure approximately 250-500 µm in diameter^9^, the relatively large contacts of commercial spinal cord stimulation electrodes, which measure about 1.3 x 3 mm^8^, are likely to activate sensory afferents across multiple adjacent DR. This non-selective activation could explain why LSCS evokes activity in both the missing limb and the residual limb.

Therefore, in this study, we designed and tested novel epidural electrode array designs with smaller contact diameters to improve the focality of the sensations evoked by LSCS. We hypothesized that using electrodes with smaller contact diameters would lead to a more precise activation of nerves that innervate distinct areas of the limb. We envisioned that these smaller contacts would generate a focused electric field, significantly less diffuse than those produced by commercial electrodes, resulting in improved selectivity. To further explore this idea, we also compared the effects of different stimulation configurations on selectivity. Specifically, we posited that bipolar stimulation, using a pair of electrodes approximately aligned with the orientation of the dorsal rootlets, would result in more selective activation of neurons projecting from the hindlimb. In contrast, we anticipated that monopolar stimulation, with a distant return electrode, would activate a larger volume of tissue.

We used a combination of animal experiments and computational modeling to examine the effects of electrode size and configuration on the selectivity of LSCS. Feline models provided neurophysiological assessments of the effects of the stimulation parameters, whereas computational models enabled us to explore a broader range of electrode designs than we could evaluate in vivo due to high fabrication costs, the complexity of surgical procedures, and the time-intensive nature of the experiments. Specifically, we simulated multipolar stimulation configurations and a broader range of electrode sizes to better understand their effects on selectivity. Together, these approaches allowed us to evaluate the trade-offs between electrode size and stimulation configuration, guiding the optimization of electrode design for the clinical translation of somatosensory neuroprosthesis for lower-limb amputees.

## Methods

We conducted acute experiments in six cats (four males; two females, Cats A – F) under approval from the University of Pittsburgh Institutional Animal Care and Use Committee. Animals were anesthetized using a ketamine/acepromazine cocktail and maintained with isoflurane (1%–2%) for the duration of each experiment. We regulated body temperature with a heating pad and monitored vital signs (heart rate, blood pressure, SpO_2_, ETCO_2_) at 15-minute intervals. Upon completion of data collection, we euthanized the animals with an IV injection of Euthasol.

### Experimental setup/surgery/implants

To quantify the activation of somatosensory afferents by LSCS, we measured antidromic evoked responses by instrumenting the sciatic and femoral trunks and their branches in the left hindlimb with nerve cuff electrodes (CorTec GmbH, Freiburg, Germany; Figure 1c, d), similar to our prior studies of sacral spinal cord stimulation and lumbar dorsal root ganglion stimulation^10–12^. Five-pole nerve cuffs with an inner diameter of 3 mm were used to record electroneurograms (ENG) from the sciatic and femoral nerve trunks. The odd numbered contacts were shorted together to act as a common reference for signals recorded from the second and fourth contacts^11,13^. The second and fourth contacts were used to record the ENG signals as they conducted along each nerve. The relative latencies of those recordings were used to calculate the conduction velocities of recruited nerve fibers^10,13^. Bipolar nerve cuffs, with diameters 1, 1.5, 2 or 2.5 mm, were used to instrument the distal nerve branches of the femoral and sciatic nerves. In total, 12 nerves were instrumented across all experiments. These nerves included the sciatic trunk (Sci), femoral trunk (Fem), saphenous (Sph), sural (Sur), common peroneal (CP), tibial (Tib), tibial distal to the branches for nerves innervating the gastrocs (dTib), and muscle nerves innervating the vasti (Vst), rectus femoris (RF), biceps femoris (BF), medial gastrocnemius (MG), and lateral gastrocnemius (LG) (Figure 1c, d). In the first experiment (Cat A), the nerve innervating the BF muscle was not instrumented due to difficulty locating it during the surgical procedure. Prior to each experiment, we measured the motor thresholds for each nerve by electrically stimulating the contacts of the nerve cuffs. We gradually increased the amplitude until a visible twitch in the hindlimb was observed, confirming that the nerve cuffs were effectively in contact with the nerve. Once we established contact, we secured sutures around each nerve cuff to ensure their stability throughout the duration of the experiment.

**Figure 1:**
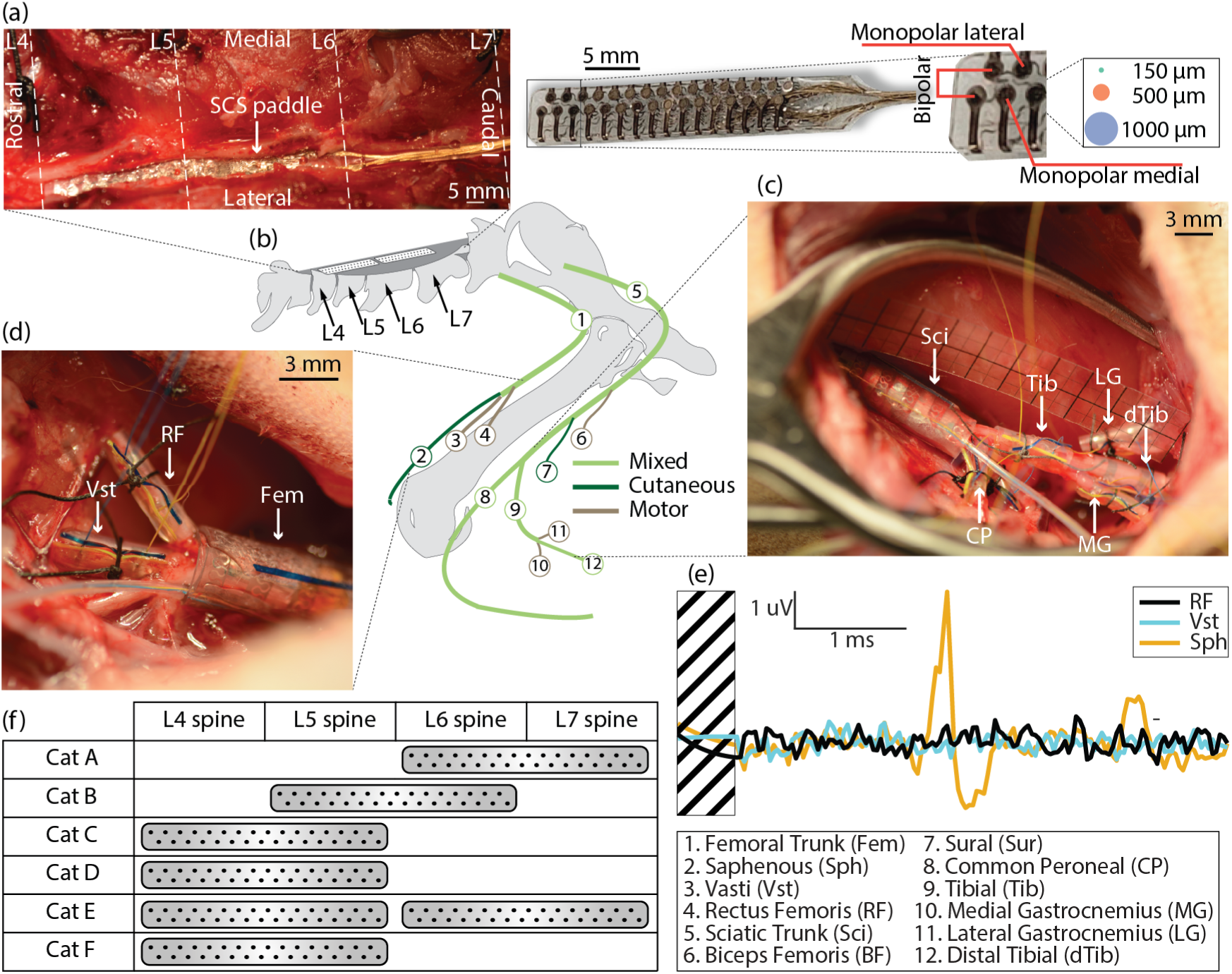
Experimental setup for LSCS and nerve cuff recordings. (a) Intraoperative image of the LSCS paddle electrode placement along the lateral surface of the spinal cord. Vertebral landmarks (L4–L7), orientation (rostral–caudal, medial–lateral), and the visible paddle contacts are labeled. (b) LSCS paddle electrodes with contact sizes of 150 µm, 500 µm, and 1000 µm. Right side of panel shows the three stimulation configurations: monopolar lateral, bipolar, and monopolar medial. (c) Dissection showing the sciatic nerve branches instrumented with nerve cuffs, including the sciatic trunk (Sci), common peroneal (CP), tibial (Tib), distal tibial (dTib), and nerves innervating the medial (MG) and lateral (LG) gastrocnemius. (d) Dissection showing the femoral nerve branches instrumented with nerve cuff electrodes, including the rectus femoris (RF), vasti (Vst), and femoral trunk (Fem). (e) Representative electroneurogram (ENG) recordings from three femoral nerve branches (RF, Vst, and Sph) showing responses to LSCS. (f) Table showing which spinal levels (L4–L7) were targeted for LSCS paddle placement in each animal (Cat 1–Cat 6) and schematic showing the 12 nerves instrumented with nerve cuffs across all experiments.

We performed a laminectomy at the L3 – S1 vertebral levels to expose the epidural surface of the lumbar spinal cord. Sutures in nearby musculature were used to mark the borders between vertebral levels prior to laminectomy. A custom 32-contact paddle electrode (CorTec GmbH, Freiburg, Germany), designed to span two vertebral levels, was placed epidurally over the lateral lumbosacral spinal cord and secured under the borders of the laminectomy. In Cats A and B, the rostral end of the paddle was placed at the rostral end of the L7 and L6 vertebrae, respectively. For subsequent experiments (Cats C, D, and F), the rostral end of the paddle was placed at the rostral end of the L4 vertebra. In Cat E, stimulation was delivered at both the L4-L5 and L6-L7 vertebral levels, with the paddle placed at the rostral end of L4 and then repositioned to the rostral end of L7. 3-cm diameter stainless steel disk electrodes were placed subcutaneously over the left and right hips to act as a reference electrode for recording and a return electrode for stimulation, respectively.

### Epidural electrode design

The paddle electrodes (Figure 1b) consisted of a platinum layer which was laser cut into a 16 × 2 array of active sites (electrode contacts). The layer of active sites was insulated with multiple layers of medical-grade silicone to ensure flexibility and durability of the paddle. Contacts in the medial column were arranged with a rostral offset so that bipolar pairs of contacts could be selected to align with the approximate orientation of the dorsal rootlets, which project at ∼45° angle with respect to the rostrocaudal (vertical) axis. The rostrocaudal (vertical) spacing between contact pairs in the same ‘row’ was 1.3 mm with a 1.6 mm mediolateral (horizontal) spacing between the contacts. There was a 2mm vertical spacing between the different rows. Three versions of the paddles were designed with all the contacts on each paddle having a diameter of either 150, 500, or 1000 µm. Across paddles, the center of each electrode contact remained in the same position, regardless of contact diameter. All experiments described below were repeated in each animal with all three electrode versions, in randomized order between animals.

### Experimental Design

We aimed to quantify the effects of electrode diameter and stimulation configuration (monopolar versus bipolar) on selectivity. By systematically analyzing these variables, we sought to understand how they influence the specific recruitment of distal nerve branches, thereby providing insights that could enhance the precision and effectiveness of LSCS. To achieve this goal, we measured the minimum LSCS current amplitude that elicited a response in any of the nerve branches (i.e., threshold amplitude) and quantified selectivity as the ability to evoke activity in one of those nerve branches to the exclusion of all other branches. While it is not possible to know what sensory percepts the animal might experience in response to selective stimulation, the underlying assumption is that activity in a single distal nerve branch would be perceived as a sensation evoked in the focal patch of skin or muscle innervated by that nerve. Additionally, we examined the range of current amplitudes over which selective recruitment was achieved (i.e., dynamic range). Finally, we classified the nerve fiber types that were activated during stimulation by measuring the conduction velocity of evoked responses in the nerve trunks.

A high amplitude survey was first conducted to determine electrode contacts that would evoke responses in one or more nerve branches ^10,12,14^. Fifty stimulation pulses were delivered at a high amplitude through each contact (or pair of contacts in the case of bipolar stimulation). This amplitude was selected through a combination of empirical and heuristic approaches to evoke responses in the nerve branches while avoiding muscle contractions and movement artifacts in the ENG signals. Across all experiments, the maximum amplitude ranged from 150 to 500 µA. Only LSCS electrode contacts that evoked responses in the nerve branches during the high amplitude survey were used in subsequent experiments to measure selectivity and dynamic range. Starting with the maximum amplitude tested, a binary search^10,12,14^ was conducted to determine the threshold amplitude for each active contact. For each stimulation amplitude during the binary search, a stimulus train of 250 symmetric biphasic pulses (cathodic-first; 66-µs pulse width; 33-µs interphase interval) was delivered through the electrode contact(s). The stimulation frequency was set based on the longest-latency neural response recorded on each nerve cuff during the high amplitude survey; we added 5 ms to this latency and took the inverse to determine the stimulation frequency, which ranged from 25 to 100 Hz. This approach allowed us to maximize the stimulation rate to minimize total experiment time. Baseline ENG activity was recorded from the nerve branches prior to each stimulation trial to be used in offline ENG analysis (see section below).

A Grapevine Neural Interface Processor (Ripple LLC, Salt Lake City, Utah) was used to deliver stimulation through a Nano 2+Stim high-current headstage. ENG activity due to stimulation was recorded using a Surf S2 headstage at a 30-kHz sampling rate. Custom MATLAB (MathWorks Inc. Natick, MA) scripts were used to modulate stimulation and record ENG^10,12^.

### Detecting Evoked Responses (ENG Analysis)

Stimulation artifacts were first removed from the recorded ENG signals by blanking a 0.5-ms window aligned with the start of each stimulus pulse and interpolating between the start and end of the blanked data^12^. This window was chosen to avoid removing responses from fast conducting fibers in any of the nerve branches while preventing contamination of ENG from the stimulus artifact. The signals were then filtered using a 2^nd^-order zero-phase high-pass Butterworth filter with a cutoff frequency at 300 Hz. For the first and fifth experiments, the femoral trunk ENGs were filtered using a 10^th^-order zero-phase high-pass Butterworth filter with a cutoff frequency at 1kHz due to large low-frequency artifacts components in the recorded ENGs, likely due to contamination from the stimulus artifact. For each stimulus train, stimulation-triggered averaging was used to remove spurious activity from individual trials and increase the signal-to-noise ratio to improve detection of CAPs^11^.

CAPs were detected from the recorded ENG signals using previously described methods^10,12,14^. For each stimulation amplitude, an ENG response distribution was determined by computing 200 stimulation-triggered averages from a random subsample of 80% of the 250 responses to individual stimulation pulses. A windowed (250-μs sliding window; 25-μs step size) root mean squared amplitude was calculated for each of the subsampled responses. Filtered baseline signals were segmented into smaller snippets, and a 99% confidence interval about the mean baseline activity was computed using the same subsampling process. If 95% of the stimulation-triggered averages within a sliding window were greater than three standard deviations above the upper bound of this confidence interval mean, the window was considered to contain a CAP.

### Selectivity and Dynamic Range

For each LSCS electrode, we first determined the threshold amplitude (i.e., the minimum stimulation amplitude that evoked a response in any of the nerve branches). At threshold amplitude, we identified distal nerve branches that were selectively recruited (i.e., the occurrence of a CAP in only one distal nerve branch, without responses in any other distal branches). If a response occurred in a distal branch and in the more proximal branching structure of the same nerve, we considered stimulation selective for only the distal branch. We then identified the dynamic range of stimulation amplitudes over which we maintained exclusive selective recruitment of that distal nerve branch. To compare the relative activation patterns of each nerve across experiments, we first organized the data according to the location of the electrode on the lumbar spine. For each instance where a nerve was selectively recruited at threshold, we identified which nerve(s) were recruited at the lowest suprathreshold that disrupted selectivity. To quantify this activation pattern for each nerve pair (nerve_i_–nerve_j_), we calculated the ratio of occurrences. The numerator was the total number of times nerve_j_ disrupted selective activation of nerve_i_, and the denominator was the total number of times nerve_i_ was selectively recruited at threshold.

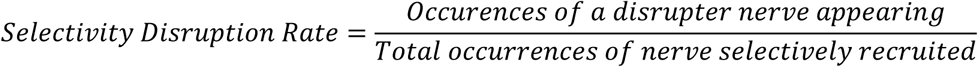

### Conduction Velocity

When CAPs were detected in the sciatic or femoral trunks, cross-correlation between signals recorded from the second and fourth contacts of the 5-pole nerve cuff was used to compute the conduction velocity (CV) of recruited afferents^10,13^. ENG signals were first normalized to a [0, 1] range and cross-correlated over a range of time delays corresponding to CVs between 10 and 125 m/s. Only negative lags, corresponding to the physiological delay where signals recorded at the fourth contact lagged those at the second, were considered. The time delay associated with the maximum cross-correlation was computed using the associated correlation lag. The CV of the evoked CAP was calculated by dividing the distance between the second and fourth contacts (8mm) by this time delay. Computed CVs were used to classify the type of recruited afferent as either Group I, Aβ, or Group II fibers.^10,13,15,16^. The estimated afferent type was used as a proxy to infer the modality of the evoked response.

### Statistics

Given that threshold amplitudes, CVs, and dynamic range followed a non-normal distribution (p<0.001, Shapiro-Wilk Test), the Kruskal-Wallis test with a post hoc Dunn’s test and a Bonferroni correction was used to evaluate pairwise differences between multiple groups of data.

Data analysis was conducted with Python Jupyter Notebooks (v7.0.8) and MATLAB (R2024a).

### Computational Modeling

We developed computational models of LSCS to examine how clinically adjustable parameters influence neural activation in the dorsal spinal roots. LSCS aims to restore selective somatosensory perception by activating large-diameter sensory afferent fibers. We therefore focused our analysis on the recruitment of Group I and Aβ fibers, which convey proprioceptive and tactile inputs critical for restoring functionally meaningful sensation.

Initially, we aimed to validate the modeling framework by comparing its predicted trends with experimental data. To accomplish this goal, we created a detailed computational model of LSCS in a feline model, positioning the electrode laterally on the dura surrounding the spinal cord. We validated the model by comparing predicted neural recruitment patterns with those observed in corresponding in-vivo experiments. After confirming that our simulations aligned with experimental observations, we extended our theoretical analysis to evaluate additional electrode configurations that we could not test experimentally. As mentioned above, it is not practical to fabricate and test every electrode design or stimulation parameter in vivo, especially when exploring novel materials or geometries. Therefore, computational models provide a scalable alternative for evaluating a wider range of electrode designs and stimulation parameters. Specifically, we simulated multipolar stimulation configurations and a broader range of electrode sizes to better understand their effects on selectivity. We also tested electrode sizes that matched the contact dimensions of a commercially available device^17^, facilitating direct comparisons between our smaller electrodes and larger commercial electrodes that are currently used in clinical settings. We constructed an anatomically realistic 3-D finite element method (FEM) model of the feline lumbar spinal cord based on imaging data from previous studies^18^. The model included spinal cord gray and white matter, dorsal and ventral roots, cerebrospinal fluid, dura mater, vertebrae, and epidural fat. To replicate our experimental conditions, we placed the custom 32-contact paddle electrode laterally on the dura aligned with the L6-L7 vertebral level (Figure 9a). The FEM model encompassed spinal roots from the L7, S1, S2 and S3 levels. To examine the influence of electrode geometry, we simulated contact diameters of 150, 500, and 1000 µm, matching those used in our animal studies. Furthermore, we also considered a contact diameter of 1.8 mm which represented the largest diameter compatible with our custom 32 contact paddle layout. Finally, we performed simulations for a single contact with an enlarged diameter of 2.5 mm to replicate the size of commercially available spinal cord stimulation paddle electrodes. This allowed us to directly compare neural selectivity achieved with our custom electrode designs to the selectivity that would be expected with existing commercial electrodes^17^.

We used previous computational modeling studies and experimental measurements to assign electrical conductivities for each tissue type^19^. We modeled each tissue as having isotropic conductivity, except for the white matter of the spinal cord and nerve roots, which we modeled as having anisotropic conductivity (Table 1).

**Table 1.** Electrical Conductivities assigned to the anatomical components in the FEM models.

| Parameter | Value (S/m) | Reference |
| --- | --- | --- |
| Gray matter | 0.230 | 39 |
| White matter (Longitudinal) | 0.600 | 39 |
| White matter (Transverse) | 0.083 | 39 |
| Dural covering | 0.600 | 40 |
| Bone | 0.020 | 41 |
| General tissue | 0.250 | 39 |
| Fat | 0.04 | 42 |

We used COMSOL Multiphysics (COMSOL, Inc., USA) to compute the extracellular potential fields generated by each LSCS configuration. For monopolar configurations, we applied a unit current (1 A) to a single electrode contact and set the outer boundaries of the model to ground (0 V), implementing Dirichlet and Neumann boundary conditions. In all simulations, we modeled the electrode paddle as a perfect insulator, and inactive contacts as equipotential with zero net current across their surface. We solved the resulting steady-state electric potential distribution by numerically solving the Laplace equation using the conjugate gradient method. We applied the principle of superposition by adding /subtracting the potential fields generated in corresponding monopolar simulations to determine the overall potential fields for bipolar and tripolar stimulation (Figure 9b).

To simulate neural responses to extracellular potential fields generated during LSCS, we implemented multicompartmental models of primary sensory neurons from the spinal roots using the NEURON simulation environment (version 8.2)^18,19^. We adapted previously published mammalian motor axon models to reflect the physiological and biophysical properties of sensory fibers. Each fiber was represented as a double-cable model, comprising nodes of Ranvier separated by three distinct myelinated segments: the myelin attachment segment (MYSA), paranodal main segment (FLUT), and internodal region (STIN)^17^. Each axon contained two concentric passive compartments, which incorporated linear leak conductances and membrane capacitance. We modeled the nodes of Ranvier with active conductances specific to sensory axons, including fast Na^+^, persistent Na^+^, fast K^+^, and slow K^+^ channels, all arranged in parallel with a linear leakage conductance and capacitance^17,20^ (Figure 9c). We randomly positioned individual axons within the spinal roots, stochastically assigning axon diameters sampled from a continuous, realistic distribution based on cadaveric measurements^21^.

We linearly interpolated extracellular potentials onto the center of each axonal compartment and applied these fields using NEURON’s extracellular mechanism via the Python interface. Because the tissue conductivities were linear, we scaled extracellular potentials proportionally with stimulation amplitude. We computed the membrane response using the backward Euler method with a 5-μs time step. We determined activation thresholds using a binary search algorithm with a resolution of 0.1 µA.

To evaluate electrode performance, we analyzed spatial selectivity as a function of electrode geometry, stimulation configuration, and amplitude. In our computational models, we calculated selectivity based on the spinal roots, as the precise innervation patterns of roots to peripheral nerves remain unclear. We computed a selectivity index (SI) for each root, r_i_, at each stimulation amplitude, I, as follows^43^:

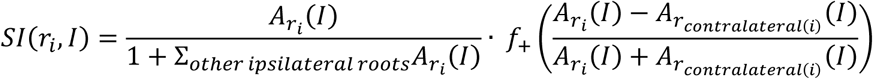

where 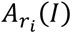 represents the percentage of fibers activated in root *r_i_*, *r_contraleral_*_(*i*)_ is the root contralateral to root *r_i_*, and 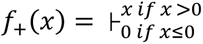. The first term of SI represents the rostrocaudal ipsilateral root selectivity. The second term measures the mediolateral selectivity and penalizes configurations that activated the root contralateral to the target rootlet^43^. If the contralateral root is activated more than the target root, then SI will equal 0. If only the target root is activated, then SI will equal 1.

## Results

Our primary goal was to design an electrode paddle capable of achieving selective activation of peripheral nerves by varying electrode diameter and stimulation configuration. To this end, we measured recruitment threshold amplitudes, evaluated selective recruitment at threshold, analyzed the dynamic range of selective recruitment, and assessed the CVs of recruited nerve fibers. We conducted these tests on six cats (A-F), using three custom paddles, each featuring a different contact diameter—150, 500, and 1000 µm—and arranged with 32 contacts in two columns. Additionally, we complemented our experiments with computational model analyses to enhance our understanding of recruitment dynamics.

### Threshold amplitudes

Recruitment threshold was defined as the minimum stimulation amplitude required to elicit a response in any of the sciatic or femoral nerve branches. Across all the experiments, recruitment thresholds ranged from 10 to 290 µA with a median amplitude of 35 µA (Figure 2). We compared the recruitment threshold amplitudes across three stimulation configurations – monopolar stimulation through the lateral columns (monopolar lateral), monopolar stimulation through the medial columns (monopolar medial) and bipolar stimulation using neighboring contacts from the medial and lateral columns (bipolar). Threshold amplitudes were significantly lower with bipolar stimulation than with monopolar medial (p=0.002) and monopolar lateral (p=0.03) stimulation (Figure 2a), though the difference was small (median thresholds for bipolar: 30 µA, monopolar lateral: 35 µA, and monopolar medial: 35 µA). No significant difference was observed in threshold amplitudes for monopolar medial vs. lateral stimulation (p>0.05). While we observed a significant difference in recruitment thresholds between electrode diameters (Figure 2b) of 500 and 1000 µm (p=0.014), the median recruitment threshold for all three electrode sizes was 35 µA. We did not observe a significant difference between recruitment thresholds of the other pairs of electrodes (150 vs. 500 µm vs: p >0.05; 150 vs. 1000 µm: p=0.19). We also investigated differences in the recruitment thresholds for stimulation across the various spinal levels. Stimulation in the L5 vertebral region had significantly lower threshold amplitudes (25 µA) compared to stimulation at the other levels (40 µA median threshold amplitudes, *p* <0.001) (Figure 2c). No other significant differences in recruitment thresholds were observed for stimulation at different vertebral levels. Lastly, we compared the recruitment thresholds across animals to assess whether there were inter-subject differences (Figure 2d). While a significant difference in threshold amplitudes was observed across all animals (p< 0.001), the thresholds in Cats A, B, and D did not differ significantly from one another, indicating a similar distribution across these three animals. In contrast, threshold amplitudes in Cats C, E, and F were each significantly different from those in A, B, and D (*p* < 0.001). Furthermore, Cats C, E, and F also differed significantly from one another.

**Figure 2:**
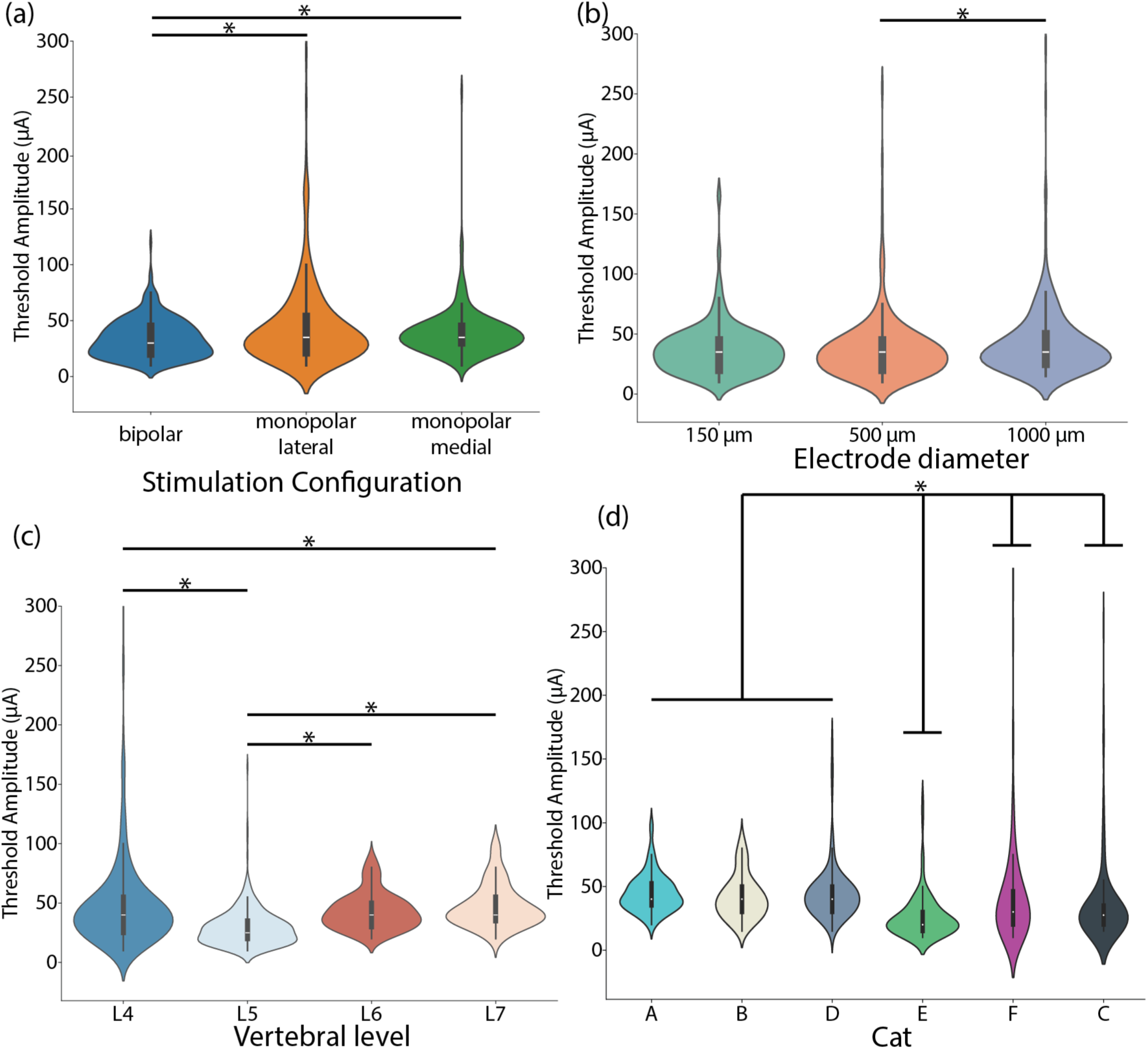
Threshold amplitudes for LSCS across stimulation parameters and experimental conditions. (a) Threshold amplitudes for three stimulation configurations: bipolar, monopolar lateral, and monopolar medial. (b) Threshold amplitudes as a function of electrode contact diameter (150 µm, 500 µm, and 1000 µm). (c) Threshold amplitudes obtained at different vertebral levels (L4, L5, L6, and L7). (d) Threshold amplitudes across individual animals. (*), p < 0.05.

### Selectivity

We define selectivity as stimulation that evoked a CAP in a single nerve branch and not in any others. When nerves in the same innervation path were activated together (e.g. tibial and distal tibial), the response was counted to be selective for the most distal nerve branch^10,14^. Across approximately 700 threshold stimulation trials, we achieved selectivity in 62% of cases across all animals. Among the three stimulation configurations, monopolar lateral was the most frequently selective (74%), followed by bipolar (62%) and monopolar medial (55%) (Figure 3a). Stimulation with 500-µm diameter electrodes achieved selectivity recruitment at threshold for 68% of LSCS electrodes, compared to 62% for both 150-µm and 1000-µm diameter electrodes (Figure 3b). Across animals, each instrumented nerve branch was selectively recruited at least once at threshold, except for the rectus femoris, which could not be recruited using monopolar medial stimulation. The distal tibial nerve was the most frequently recruited (32%) sciatic nerve branch, whereas the saphenous nerve was the most frequently recruited (16%) femoral branch. Both nerves predominantly convey cutaneous signals from the skin. Recruitment patterns for stimulation across the different lumbar regions were similar regardless of stimulation configuration (Figure 4) or electrode diameter (Figure 5). At the L4 vertebral region, femoral branches were mainly selectively recruited. Both femoral and sciatic nerve branches were recruited at the L5 vertebral region. However, monopolar medial stimulation mostly recruited sciatic nerve branches at the L4 and L5 vertebral levels. Stimulation at the L6 and L7 vertebral levels only recruited sciatic nerve branches regardless of stimulation configuration.

**Figure 3:**
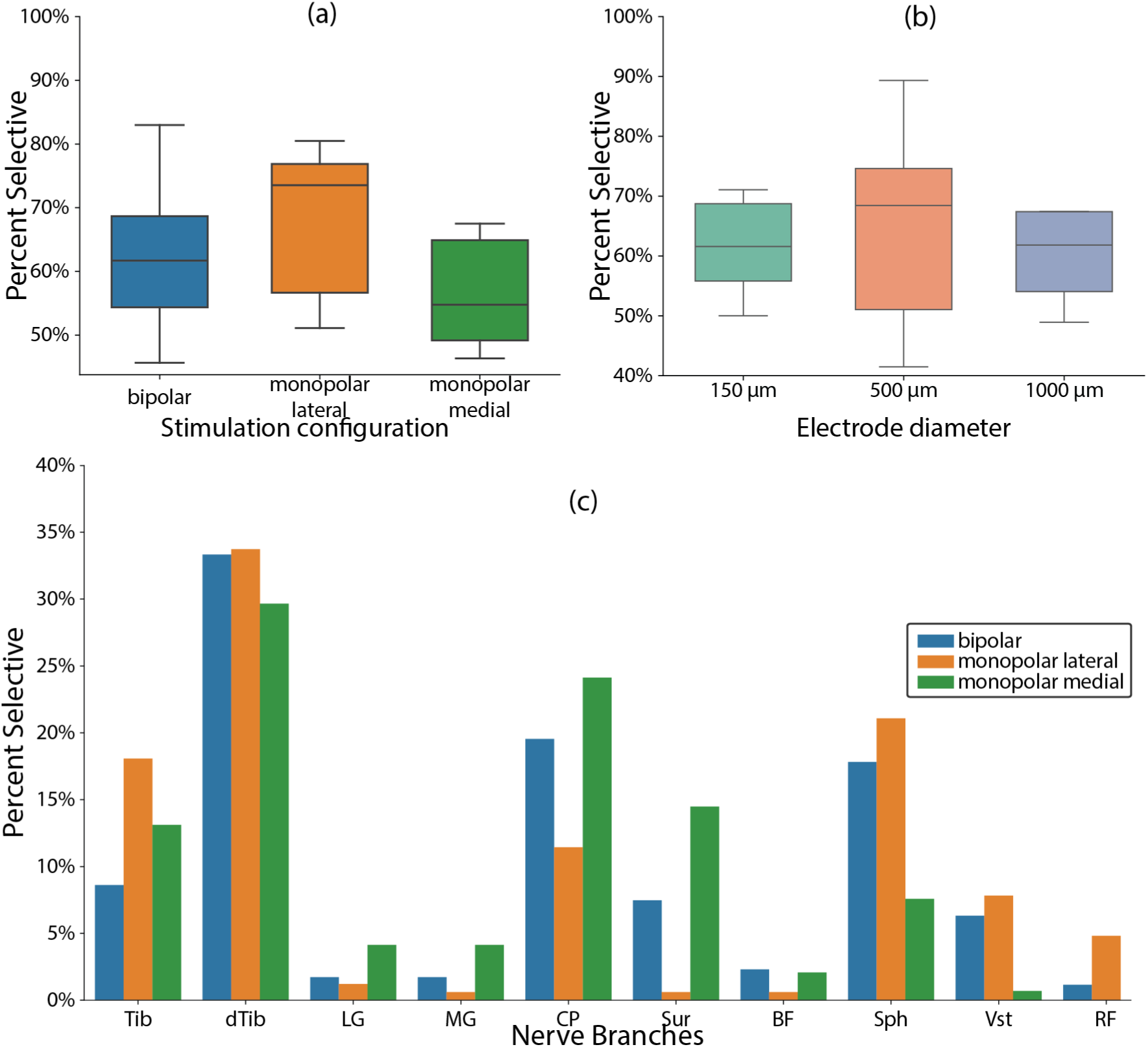
Percent selective responses across stimulation parameters and individual nerves. (a) Percent of responses that were selective to a single nerve for each stimulation configuration: bipolar, monopolar lateral, and monopolar medial. (b) Percent of selective responses across electrode diameters (150 μm, 500 μm, and 1000 μm). Box plot distributions in (a) and (b) are across different animals. (c) Percent of selective responses for each individual nerve branch, grouped by stimulation configuration. Colors correspond to stimulation types as shown in the legend.

**Figure 4:**
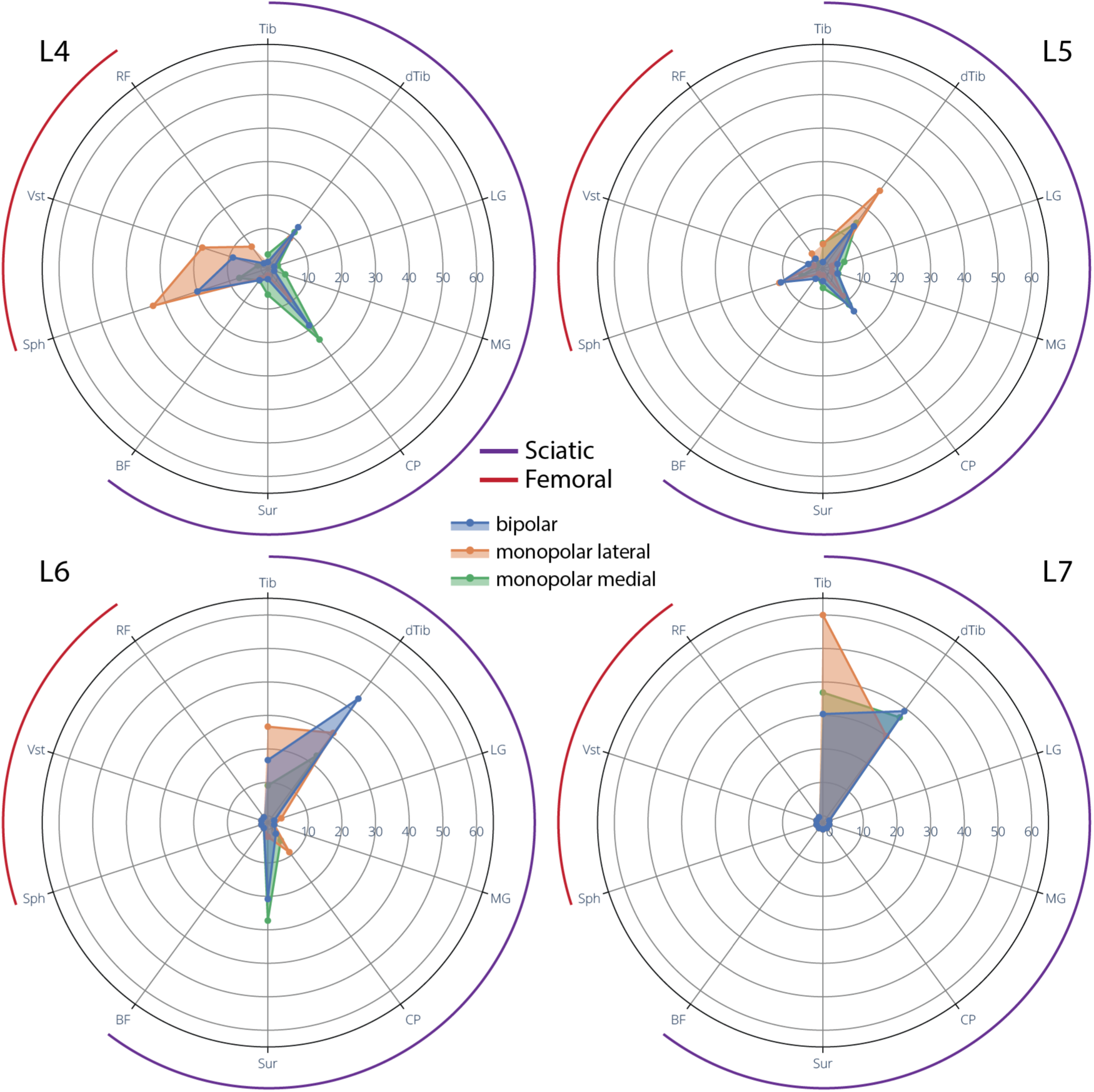
Polar plots showing selective nerve recruitment by vertebral level and stimulation configuration. Polar plots illustrate the percent of selective activations across individual femoral (black arc) and sciatic (purple arc) nerve branches during LSCS at vertebral levels L4, L5, L6, and L7. Each plot represents one vertebral level, and within each plot, shaded areas show the distribution of selective responses for three configurations: bipolar (blue), monopolar lateral (green), and monopolar medial (orange). Nerve labels are arranged circumferentially and include branches of both the femoral and sciatic nerves.

**Figure 5:**
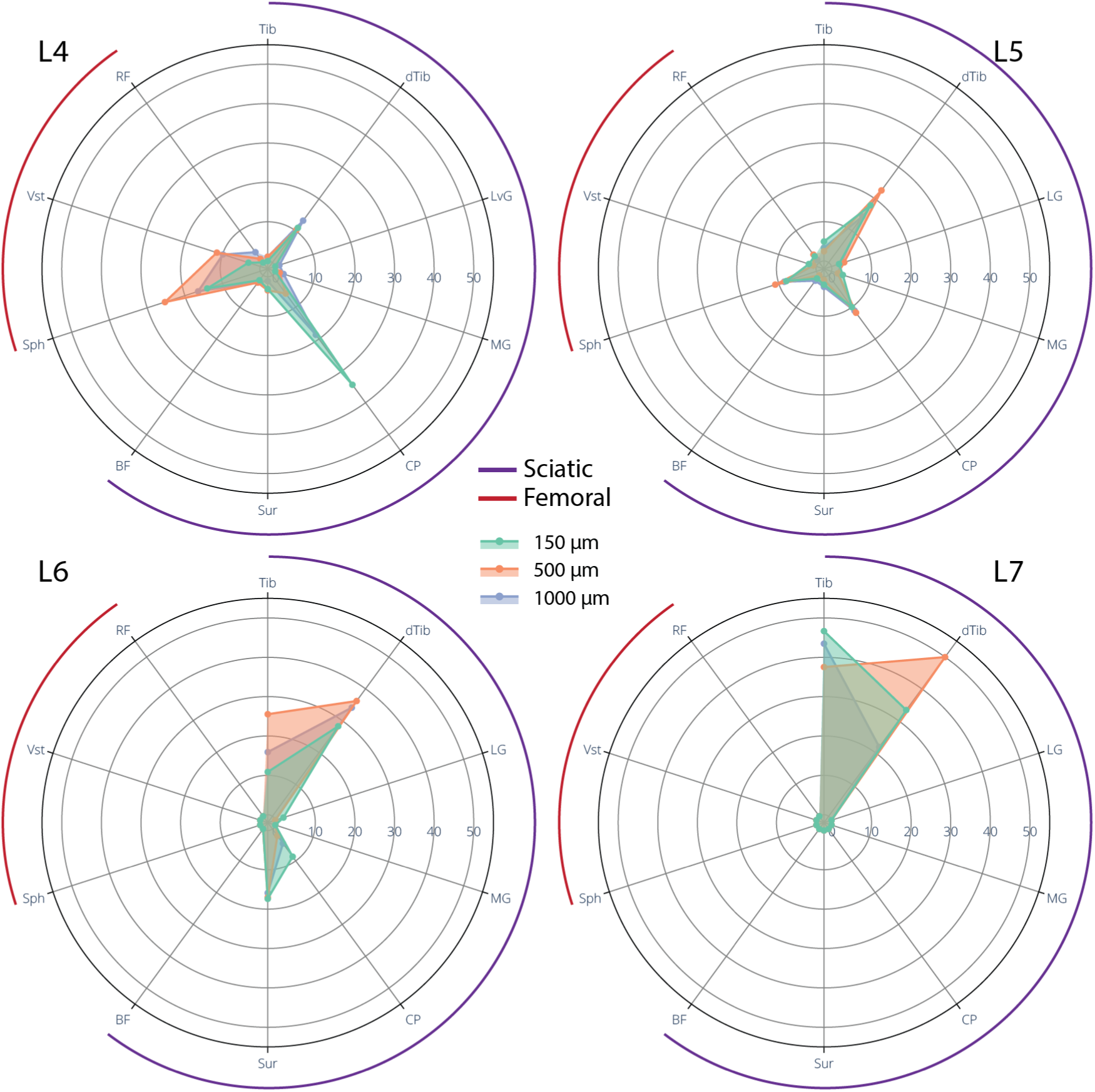
Polar plots showing selective nerve recruitment by vertebral level and electrode size. Polar plots illustrate the percent of selective activations across femoral (black arc) and sciatic (purple arc) nerve branches during LSCS at vertebral levels L4, L5, L6, and L7. Each plot corresponds to one vertebral level, and shaded areas represent selective responses for different electrode size: 1000 µm (blue), 500 µm (orange), and 150 µm (green).

### Dynamic Range

Dynamic ranges were significantly higher for monopolar lateral stimulation compared to monopolar medial stimulation (p < 0.001), although the difference was very small and there was no difference between monopolar and bipolar stimulation (median dynamic range for monopolar lateral, monopolar medial, and bipolar were all 5 µA) (Figure 6a). Statistically significant differences were observed in the dynamic ranges across lumbar spine regions (Figure 6b), with stimulation in the L4 region demonstrating a higher dynamic range compared to stimulation in L5 (p<0.001), L6 (p=0.007), and L7 (p < 0.001). There was also a significant difference observed between L5 and L6 (p=0.007), with L6 showing a higher dynamic range than L5. No other significant differences were found in the dynamic range of stimulation across the lumbar spine. Again, however, these differences were small (less than 10 µA). There were no significant differences found between the dynamic ranges of the three electrode diameters.

**Figure 6:**
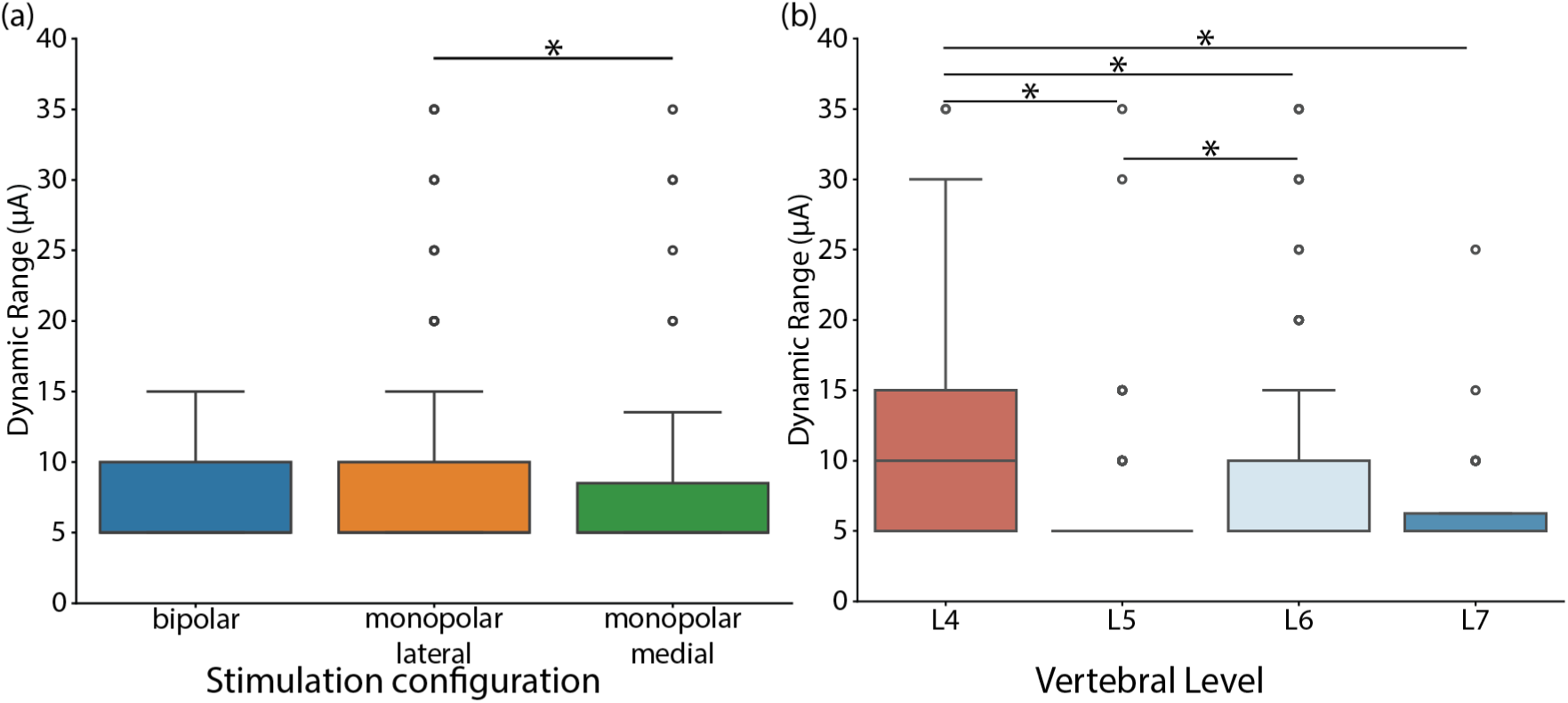
Dynamic range of stimulation across configurations, vertebral levels, and electrode sizes. (a) Dynamic range of stimulation (defined as the range of current amplitudes over which selective recruitment was achieved) for each stimulation configuration: bipolar, monopolar lateral, and monopolar medial. (b) Dynamic range at different vertebral levels (L4, L5, L6, L7).

We also investigated nerve co-activation at the lowest amplitude above threshold where another nerve was recruited (Figure 7). Our results showed that disruption of selectivity was often due to recruitment of nerves branches belonging to the same nerve trunk (i.e., sciatic or femoral). However, this pattern was most prevalent with monopolar lateral stimulation, with monopolar medial and bipolar stimulation more frequently eliciting cross-trunk coactivation.

**Figure 7:**
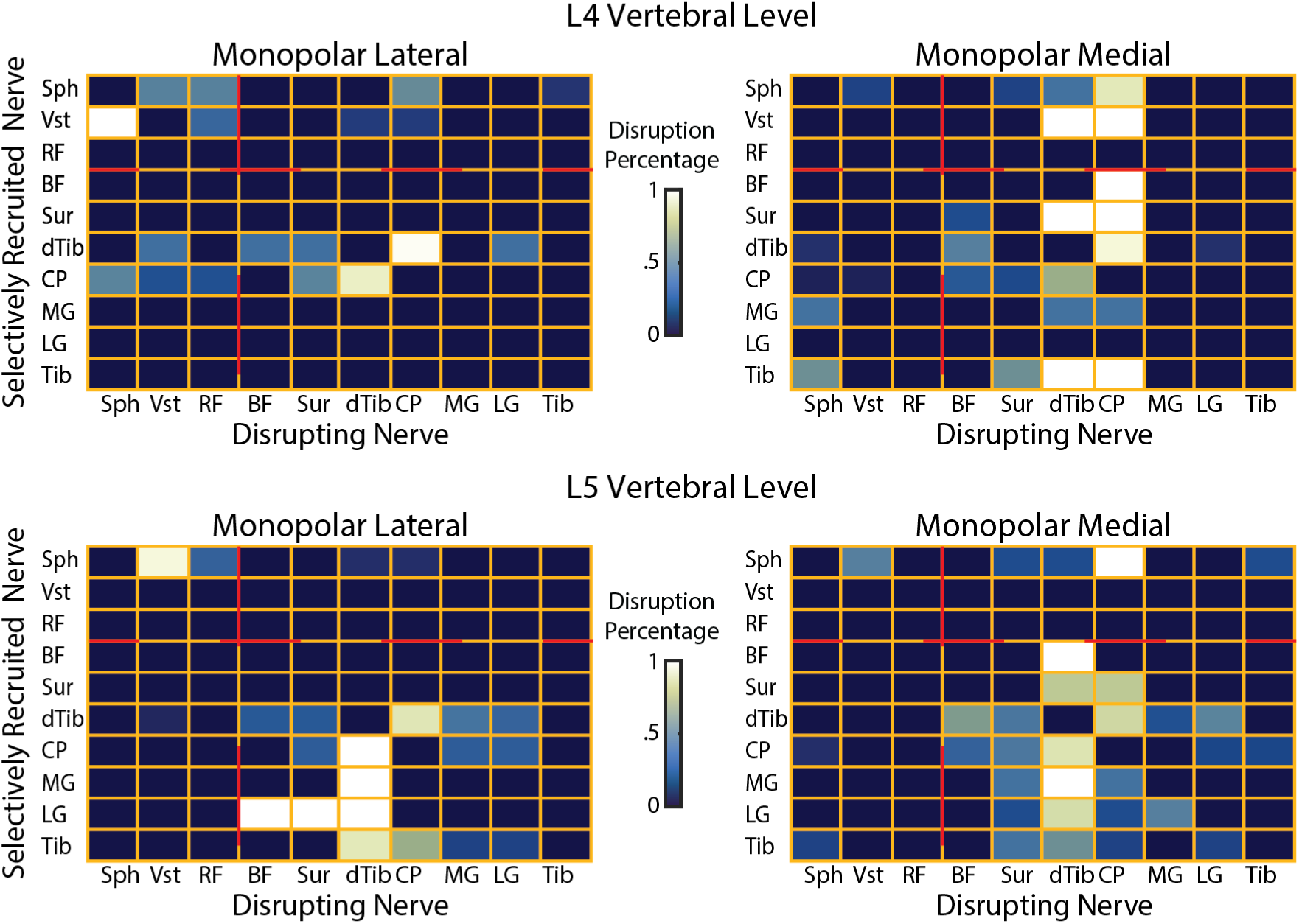
Co-activation patterns during monopolar stimulation across vertebral levels. Heatmaps showing loss of selectivity patterns during monopolar lateral and medial stimulation at vertebral levels L4 and L5. Rows represent nerves selectively recruited at threshold, and columns represent nerves that were subsequently co-activated and disrupted selectivity. The color of each cell reflects the ratio of co-activation events between each nerve pair. Red dashed lines separate sciatic and femoral nerve branches.

### Conduction Velocity

The conduction velocities of the recruited afferents in the sciatic and femoral trunks were compared at threshold amplitude to identify their fiber type (Figure 8). Here, we only considered responses in the femoral and sciatic trunks at the lowest amplitude that evoked a response in either nerve. In the sciatic nerve trunk, we saw activation of either Group I (80-120 m/s) or Aβ (60-80 m/s) fibers, while in the femoral trunk, we activated Group I (80-120 m/s), a mixture of Aβ /Group II (40-80 m/s), and very few slow conducting Group II only fibers (<35 m/s). Slow-conducting fibers in the femoral trunk were recruited only using bipolar electrodes (Figure 8b).

**Figure 8:**
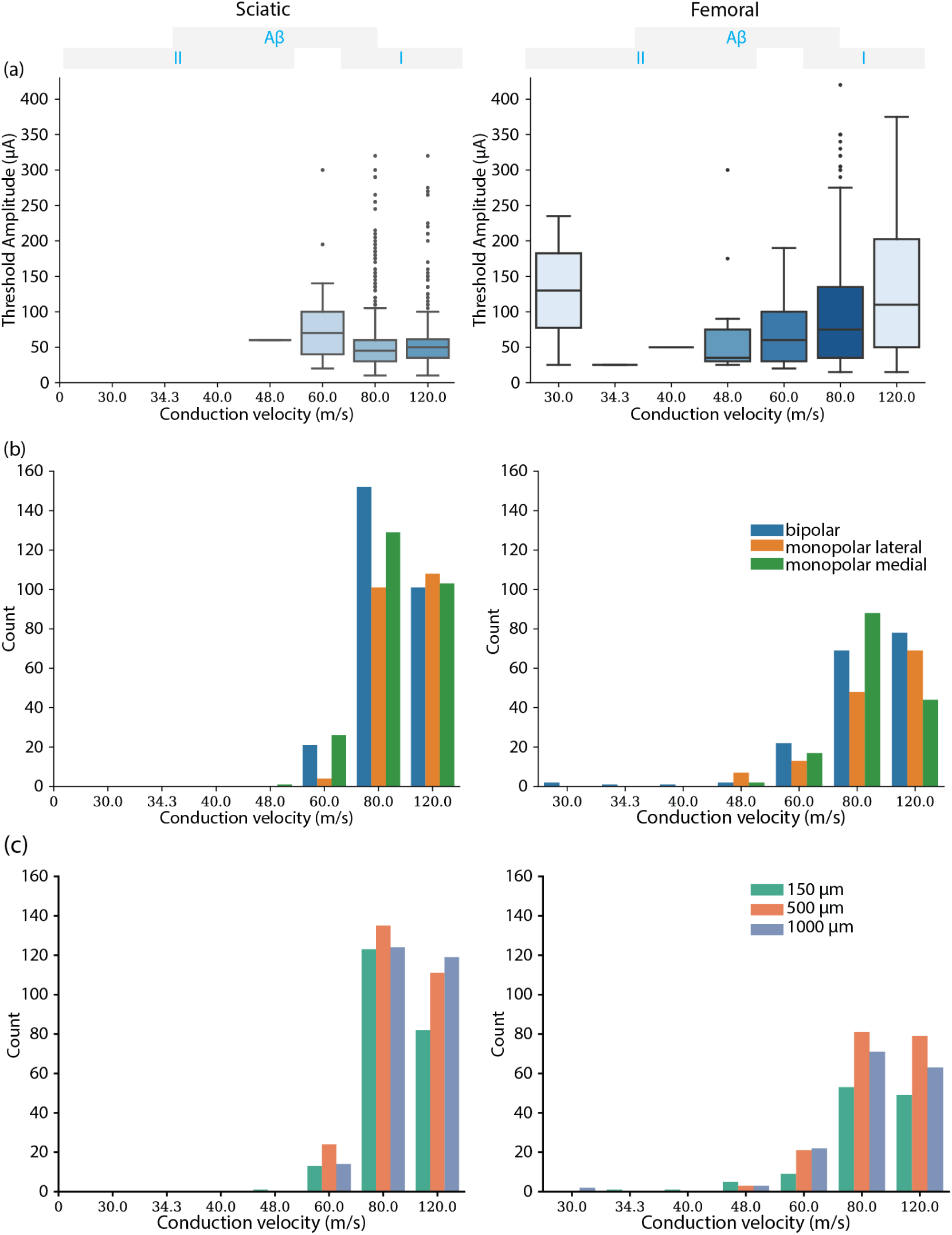
Conduction velocity distribution and corresponding thresholds across sciatic and femoral nerves. (a) Threshold amplitudes as a function of CV. (b) Distribution of recruited fiber CVs across different stimulation configurations: bipolar, monopolar lateral, and monopolar medial. (c) Distribution of recruited fiber CVs across different electrode contact diameters (0.15, 0.5, and 1.0 mm). CV values were discretized due to sampling frequency limitations of the recording setup.

### Computational Model

Our computational models investigated electrode contact diameters of 150, 500, and 1000 µm, positioned laterally at the L6-L7 vertebral level (Figure 9a, b). The models revealed minimal differences in selectivity and threshold values for target spinal roots among these electrode sizes (Figure 9d, e), findings that were consistent with our experimental data. Although we included axons at the rootlet levels in the model, the stimulation was unable to selectively recruit individual rootlets. Instead, it recruited axons from multiple adjacent rootlets simultaneously, regardless of electrode diameter.

**Figure 9:**
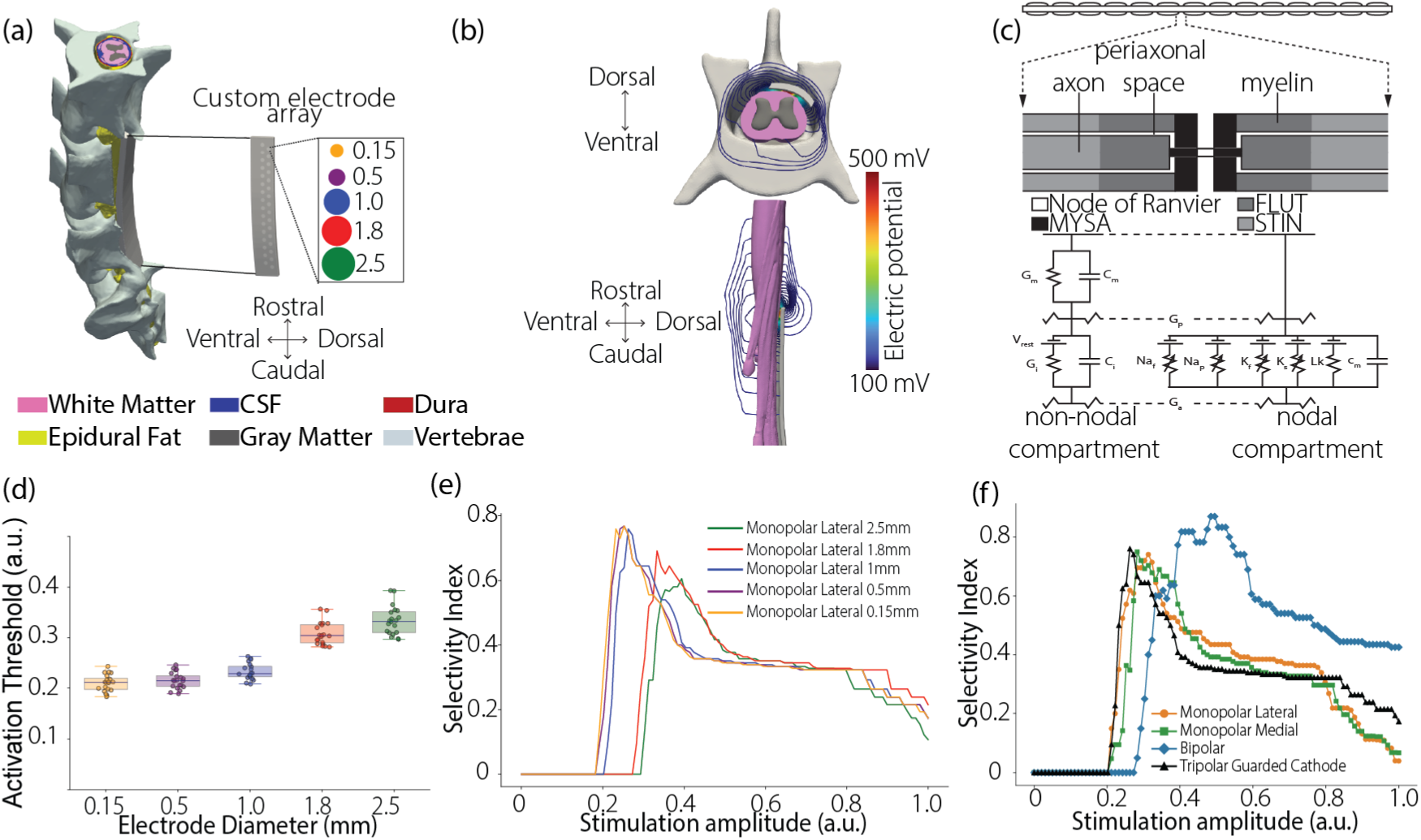
Computational modeling framework for lateral spinal cord stimulation. (a) Canonical model of the lumbosacral feline spinal cord encompassing spinal levels L5 through L7. (b) Spatial distribution of extracellular voltage generated by the custom paddle electrodes. (c) Multicompartment cable models of primary sensory neurons, representing the biophysical channel dynamics of large diameter myelinated Aα and Aβ fibers. This panel was adapted from reference 42. (d) Dependence of neural activation thresholds on electrode size. (e) Computed selectivity index as a function of electrode size, reflecting preferential activation of target fibers. (f) Computed selectivity index as a function of stimulation configuration.

The computational models also enabled us to simulate large electrode contacts, including a 1.8-mm diameter, which represented the largest individual contact size possible within our custom array layout while still accommodating the full set of 32 electrodes. Additionally, we simulated a single 2.5-mm diameter electrode, corresponding to the typical surface area of individual contacts used in commercial SCS systems. It is important to note that the 1.8-mm size was constrained by our array’s physical configuration, which required fitting all 32 electrodes within the paddle dimensions. In contrast, the 2.5-mm configuration modeled only a single electrode, rather than a full multi-electrode array. With these larger sizes, we observed an increase in recruitment threshold and a decrease in selectivity (Figure 9d, e). We also simulated a tripolar guarded cathode configuration, in which three consecutive contacts along either the medial or lateral line of contacts were used. The middle contact served as the cathode (active), while the two flanking contacts acted as the return (anodes). Although this configuration was not universally the most selective, it activated neurons at relatively lower stimulation amplitudes compared to monopolar and bipolar configurations (Figure 9f).

## Discussion

In this study, we assessed the effect of electrode diameter and stimulation configuration on selectivity of epidural LSCS. Through a combination of in-vivo and in-silico models, we aimed to assess how specific design parameters influence neural recruitment during LSCS. In our in-vivo experiments, we compared the effects of LSCS using three contact diameters – 150, 500, and 1000 µm – and three stimulation configurations – monopolar lateral, monopolar medial, and bipolar. Our computational models allowed us to test additional parameters that were not feasible to test in-vivo. Specifically, we simulated a 1.8-mm contact diameter, the largest size possible while maintaining a similar contact pattern design as the initial configurations and still fitting within our custom paddle. We also tested a single 2.5-mm diameter contact, which matches the size of those used in commercially available spinal cord stimulation electrodes, to allow for direct performance comparisons. We positioned this contact at a similar rostrocaudal level as the rostral-most electrode pair of our custom paddle to enable fair comparison. To enhance stimulation selectivity, clinicians often use a tripolar guarded cathode configuration in SCS, where three neighboring contacts along the rostrocaudal axis are used as a central cathode with surrounding anodes^20–22^. Therefore, we also assessed the effects of this stimulation configuration on neural selectivity in LSCS.

In our experimental study, we found that lateral epidural stimulation could successfully achieve selective activation of nerve branches across the hindlimb at relatively low stimulation amplitudes. The observed activation patterns corresponded well with expected dermatomes, indicating a level of precision in targeting specific nerve pathways. Despite this promising selectivity, the dynamic range was consistently narrow, suggesting there may be inherent limits on selectivity of epidural LSCS. When we tested different electrode contact sizes and configurations, the effects on selectivity were minimal, resulting in changes that may be functionally negligible. This outcome hints that alterations in physical configuration alone might not lead to significantly enhanced selectivity of LSCS. These findings lead us to consider the possibility of fundamental biological constraints inherent in the organization of the dorsal rootlets^23^. Such structural limitations could inherently cap the degree of selectivity achievable with LSCS. This complexity in dorsal rootlet arrangement may mean that even with advanced electrode design optimizations, there is a ceiling to the precision that LSCS can offer.

Our modeling study demonstrated that there are benefits in reducing contact size below those used in existing clinical SCS systems. Additionally, the computational models provided insights that helped explain why modifications in electrode dimensions within the range tested in experimentally did not translate to markedly improved selectivity. Specifically, we found that electrodes with diameters above 1000 µm activated neurons projecting from multiple neighboring spinal roots, while those with diameters of 1000 µm or less activated neurons projecting from multiple rootlets within a single spinal root. As such, our findings also demonstrate that, after a certain point, making contacts smaller yields rapidly diminishing returns in terms of improving selectivity, because the high conductivity of the CSF shunts currents, thereby reducing specificity of smaller contacts. The modeling study underscores the importance of optimizing electrode design within these parameters, as increasing contact size compromises the focused activation of specific nerve pathways. Therefore, our findings advocate for a strategic approach that respects these optimal size limits while exploring other avenues, such as innovative configurations or stimulation techniques, to further improve the selectivity achievable with LSCS. This holistic approach is crucial for advancing LSCS methodologies and enhancing their clinical application for precise sensory restoration in patients.

With the underlying hypothesis that individual dorsal rootlets carry specific nerve information, we anticipated that electrodes with smaller contact diameters would exhibit greater selectivity during LSCS. We hypothesized that the smaller electrodes would enable us to activate specific nerve pathways more effectively, thus enhancing the precision and focality of the evoked sensations. However, we did not observe substantial differences in selectivity across the contact diameters that were tested in-vivo. The recruitment thresholds for all the contact diameters had similar median values and thresholds were usually low (∼25-50 µA). These results were consistent with our findings in-silico. In our computational models, we observed that the 150 µm, 500 µm, and 1000 µm electrodes exhibited similar recruitment thresholds and selectivity, with no significant differences among the three contact diameters. Although we included axons separated at the rootlet level in our models, the electrodes primarily recruited whole spinal roots, so we did not observe rootlet-level selectivity. These findings suggest that even smaller electrodes lack the precision to target specific rootlets and instead engage neural elements across a broader area at the spinal root level^23^. The electric field distribution patterns remained consistent across all the three sizes, and the electrodes were small enough to avoid activating axons from non-target levels. However, when we simulated electrode contact diameters of 1.8 mm and 2.5 mm, we observed a decrease in target spinal root selectivity due to the recruitment of axons from non-target spinal roots. The wider electric field spread by the larger electrodes easily recruited axons from the contralateral spinal cord, further reducing selectivity at the root level. Additionally, these larger electrodes increased the activation threshold and recruited fibers at a higher amplitude due to their lower impedances from increased surface area.

During our in-vivo testing of different stimulation configurations, we anticipated greater selectivity with bipolar stimulation due to the arrangement of the bipole in the custom paddle^24^. This orientation aligned the pair of electrodes approximately parallel to the dorsal rootlets, directing the current more precisely from source to sink. Unlike monopolar stimulation, where the distant sink results in a broader activation of tissue, in the bipolar setup, a substantial portion of the current flows directly between the working electrode and the return electrode, with extracellular voltage theoretically dropping with the square of the distance from the electrode, rather than linear as with monopolar stimulation^22^. In our experiments, we found no significant differences in selectivity among the three stimulation configurations tested. While stimulation thresholds were generally comparable, we noted that bipolar stimulation had a statistically lower recruitment threshold than either medial or lateral monopolar stimulation (Figure 2a).

To further investigate the effects of stimulation configuration on selectivity, we turned to a computational model to test a tripolar guarded cathode configuration, known for its clinical application in spinal cord stimulation. Tripolar configurations provide a focused and directed current flow, which can minimize the activation of unintended neural targets and reduce side effects. By adopting this setup, we aimed to explore whether the tripolar configuration could improve selectivity. Through our models, our goal was to determine if this configuration could address some of the limitations observed with other setups. We found that there was no significant difference in selectivity across thresholds among the three configurations tested, although the bipolar configuration was slightly more selective likely because the associated contacts were aligned along the dorsal rootles. In our computational models, although the tripolar configuration was not the most selective, it achieved recruitment at relatively lower thresholds compared to monopolar and bipolar configurations due to its highly directed current flow. This outcome suggests that while this configuration may not offer superior selectivity, it can efficiently recruit neurons with less energy, potentially enhancing comfort and effectiveness for clinical SCS applications. It is also important to note that the selectivity and efficiency of the tripolar configuration largely depend on the relative orientation of the electrodes to the target neurons. If the neural trajectories were better aligned with the direction of the current flow, the tripolar configuration would likely demonstrate greater selectivity, which is why they are highly valuable in traditional SCS where the dorsal column axons are aligned with the orientation of the longitudinal guarded cathode. In our models, we did not see a significant difference because the bipolar contacts were already aligned with the nerve trajectories, minimizing the benefits typically associated with tripolar guarded cathode configurations.

Differences in recruitment patterns were most evident when the different stimulation parameters (contact diameter and configuration) were compared across the different lumbar levels. Consistent with reported tactile dermatomes in cats^25,26^, we observed that stimulation in the L6 and L7 vertebral level selectively recruited sciatic nerve branches that innervate the foot, regardless of stimulation configuration or electrode contact diameter. This trend supports results from our previous study^10^ that suggest that stimulation in this region may be most relevant for clinical applications (e.g., restoring sensory feedback from the missing foot of people with a lower-limb amputation). Likewise, monopolar lateral stimulation in the L4 region resulted in expected selective recruitment patterns of femoral nerve branches as compared to bipolar stimulation in the same region. Monopolar medial stimulation in the same region recruited mostly sciatic nerve branches. This may be due to the proximity of the electrodes to the dorsal columns, which allowed for activation of fibers from the sciatic nerve that travel rostrally in the dorsal columns towards supra-spinal structures of the nervous system. A mixture of sciatic and femoral nerve branches was selectively recruited for stimulation in the L5 spine.

Across all stimulation parameters, the dynamic range was consistently small, typically around 5 – 10 µA. This range means we moved from selective to non-selective responses with a single step-size increase in stimulation amplitude (for our given hardware constraints). This phenomenon likely reflects the presence of neurons projecting from multiple peripheral nerves in a single dorsal rootlet. However, we observed a clear somatotopic organization across spinal levels, where co-activated nerves often originated from the same nerve trunk (femoral or sciatic). This trend was most prevalent during monopolar lateral stimulation. This again may be because medial and bipolar stimulation tend to activate neurons in the dorsal columns that project from multiple dorsal roots.

Stimulation at threshold primarily activated fibers with conduction velocities ranging from 60 to 120 m/s, corresponding to medium- and large-diameter myelinated fibers within the sciatic and femoral nerve trunks. Although we do not report the recruitment pattern at supra-threshold stimulation amplitudes, the activation of fast-conducting fibers at low amplitudes is consistent with previous studies that show a large-to-small diameter progression of fiber recruitment as stimulation amplitude increases^27^. The recruitment of large-diameter fibers at threshold amplitude is particularly relevant in the clinical context of somatosensory restoration for people with lower-limb amputations. Given that 80-85% of lower-limb amputees experience phantom limb pain^28,29^, the activation of these fibers via epidural spinal cord stimulation offers a promising avenue to engage spinal-gating mechanisms that attenuate nociceptive sensations originating from the phantom limb^30–33^. Epidural LSCS may provide a therapeutic benefit by possibly alleviating phantom limb pain in people with a lower limb amputation.

Results from our in-vivo and computational experiments underscore the role of electrode geometry and stimulation configuration in neural recruitment dynamics. Epidural LSCS achieved selective activation of nerve branches throughout the hindlimb at low stimulation amplitudes, and the patterns of activation matched expected dermatomes^17^. However, contrary to our assumption that smaller electrodes will inherently offer better selectivity, our findings suggest that reducing electrode dimensions beyond a certain threshold does not provide any advantages. Instead, it compromises the benefits by increasing contact impedance, which hampers effective stimulation in the long run. As electrodes become smaller, the increased impedance requires higher power consumption to maintain efficient current delivery. This heightened power demand strains the energy efficiency of the device and raises the risk of tissue damage. Excessive power can potentially damage adjacent tissues and compromise the safety and longevity of the electrode-tissue interface^34^. Taken together, these results strongly suggest that there may be fundamental neuroanatomical limitations of epidural LSCS. We explored this question further in a companion paper^23^ in which we present results of a similar study in the same set of animals, measuring the responses to stimulation of individual dorsal rootlets. In that study, we did, in fact, find that multiple rootlets within a given spinal level included sensory afferents projecting from broad and overlapping regions of the limb^23^.

In the complex system of epidural stimulation, electrodes are placed epidurally, typically surrounded by fat. The electrical current must traverse the dura matter before entering the cerebrospinal fluid (CSF), a highly conductive medium surrounding the target white matter. The relatively high conductivity of the CSF causes substantial shunting of the current. This effect undermines target specificity and induces undesired activation of non-target areas, limiting the ability to precisely stimulate the intended white matter regions and diminishing the overall efficacy of the stimulation. Our models re-emphasize the importance of considering both electrode size and the biophysical environment in which the current propagates. Achieving optimal selectivity in such systems requires balancing electrode dimensions with power efficiency and tissue safety.

Additionally, results from varying stimulation configurations indicate that there may not be an extra functional benefit in multipolar configurations, although the bipolar and guarded cathode configurations had lower stimulation thresholds, they did not offer an improved selectivity over using lateral monopolar electrodes. The consistently small dynamic range of selectivity suggests it may be challenging to find selective stimulation parameters. We believe there may be fundamental limitations in the organization of the dorsal rootlets that set a ceiling for the selectivity that can be achieved with LSCS.

### Limitations

While this study presents a promising avenue to characterize the selective recruitment patterns of epidural LSCS using our custom electrodes, translation to humans will require additional work. Our results showed that there may be a benefit in decreasing electrode size from existing clinical SCS devices, although only to a certain limit, but future studies will be required to confirm those results in humans. Further, given that our study was conducted in anesthetized cats, we cannot report the limb location and qualitative nature of any evoked sensation. While our studies present the potential for these paddle electrodes to be used for somatosensory neuroprosthetic applications in lower-limb amputees, testing in humans will be required to quantify the location and quality of the evoked percepts. We used CV as a proxy for the modality of the sensations that would be evoked by stimulation, although there is overlap in the conduction velocities of secondary muscle spindles and Aβ tactile afferents. Further, across most recent studies of clinical somatosensory neuroprosthesis it has been difficult to evoke proprioceptive sensations, even though primary muscle spindles are the largest afferents with the lowest thresholds, suggesting that it may not be sufficient to simply activate these neurons to evoke percepts. An additional limitation of our study design is that the data acquisition system has a sampling frequency of 30kHz, which limited our ability to differentiate conduction velocities, with a coarser resolution for the fastest conduction velocities. Therefore, the reported velocities are possibly approximations of the actual conduction velocities of the recruited afferents.

For each animal, we assumed that we placed different diameter electrode paddles in the same anatomical position. We used sutures near the spinal cord to align the paddles within an animal. Although this alignment was coarse and minor misalignments might have influenced differences in results across paddles with various contact diameters, the consistency of the results across diameters and cats suggests that this was not a significant limitation of the experiment. Furthermore, while our results demonstrate the electrodes’ ability to selectively recruit nerve branches, we did not evaluate their long-term safety when stimulating neural tissue. Future studies should conduct chronic experiments to assess the long-term safety of these electrodes. These studies should also quantify potential lead migration after electrode implantation, as this remains a common issue with human spinal cord stimulation implants.^35–38^.

## Conclusion

In summary, our study demonstrates that the use of novel high-density flexible paddles with smaller electrode sizes enables the selective recruitment of sciatic and femoral nerve branches. However, the small dynamic range of selectivity is inconsistent with our hypothesis about the functional organization of the dorsal rootles and their projections into the nerve branches of the foot and leg. While our results highlight the potential of these electrodes to be used in providing focal sensations, future studies should investigate the functional organization of the dorsal rootlets as that may inform design choices for these novel electrodes. Here, we propose a pathway toward designing electrodes specifically tailored for somatosensory feedback restoration in individuals with lower-limb amputations, addressing both sensory deficits and therapeutic needs. Epidural LSCS achieved selective activation of nerve branches throughout the hindlimb at low stimulation amplitudes, and the patterns of activation matched expected dermatomes^17^. However, the dynamic range of selectivity was always small, suggesting it may be challenging to find selective stimulation parameters. Different contact sizes and configurations only had minor effects on selectivity, likely at a level that has minimal functional impact. There may be fundamental limitations in the organization of the dorsal rootlets that set a ceiling for the selectivity that can be achieved with LSCS^23^.

## Supplementary Material

**Supplementary Figure 1:**
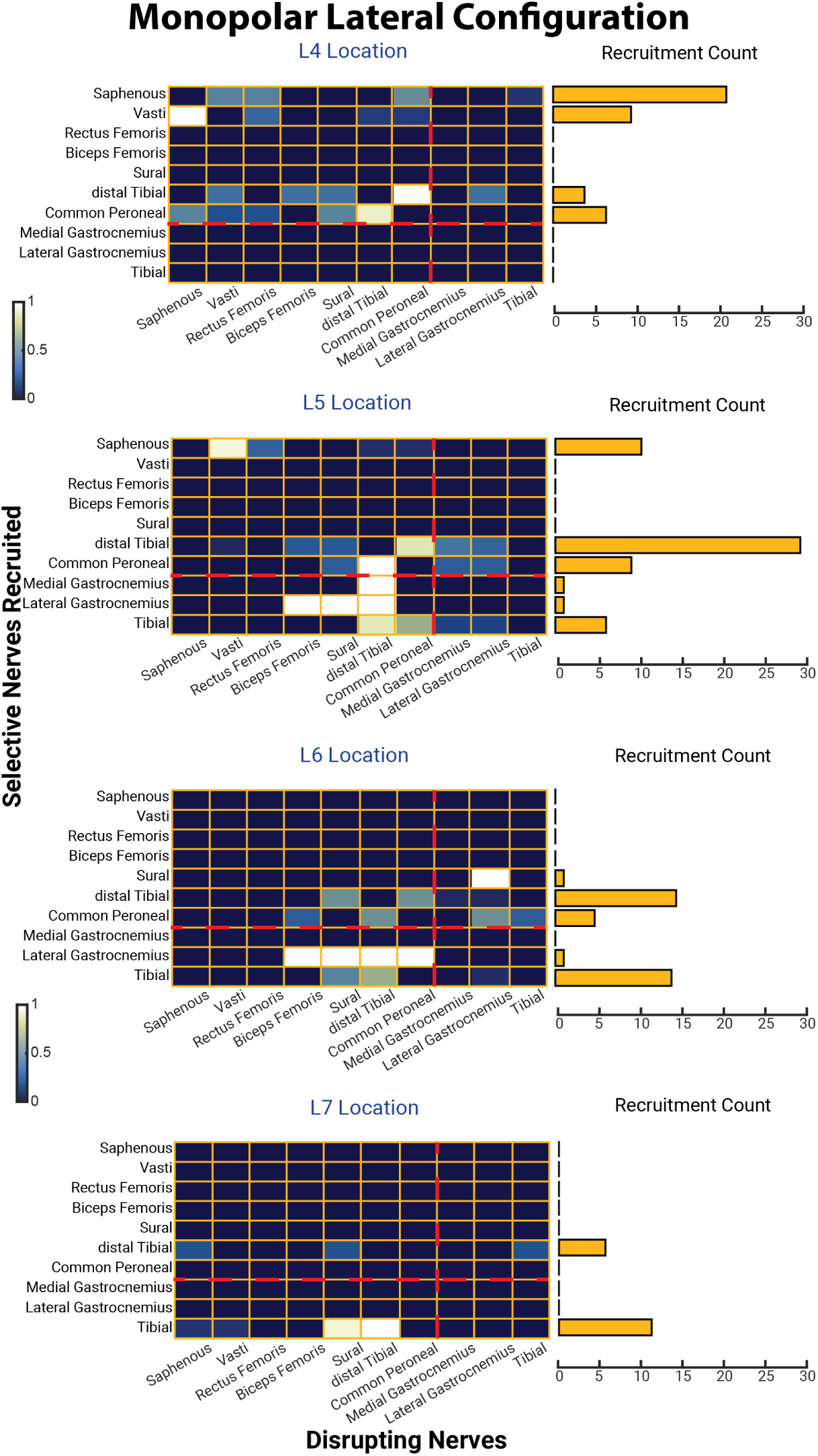
Co-activation patterns during monopolar lateral stimulation across vertebral levels. Heatmaps show co-activation patterns at the lowest amplitude where loss of selectivity occurred during monopolar lateral stimulation at vertebral levels L4, L5, L6, and L7. Rows represent nerves selectively recruited at threshold, and columns represent nerves that were subsequently co-activated and disrupted selectivity. The color of each cell reflects the ratio of co-activation events between each nerve pair. Red dashed lines separate sciatic and femoral nerve branches. Adjacent bar plots indicate how often each nerve was involved in recruitment loss as a disrupter nerve.

**Supplementary Figure 2:**
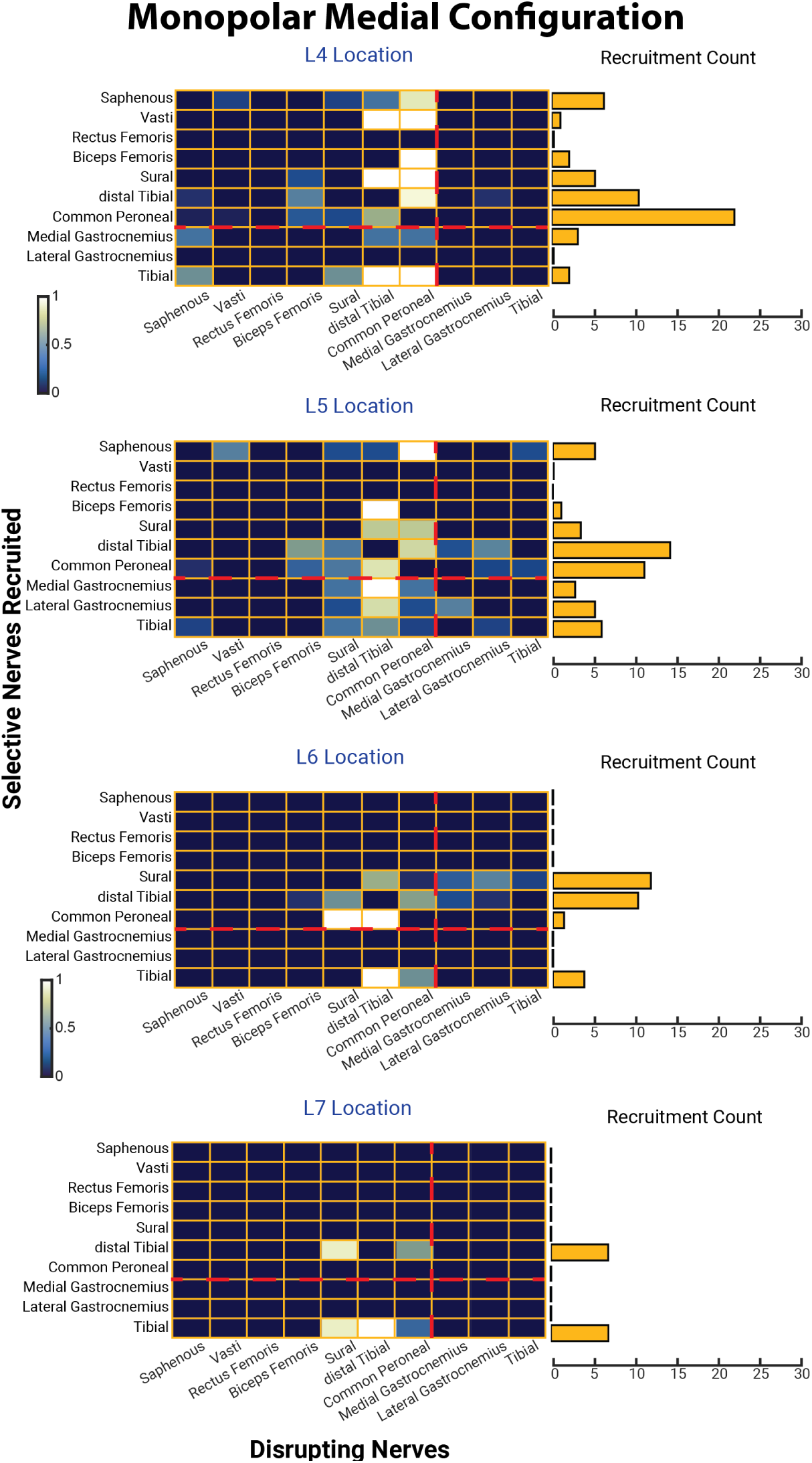
Co-activation patterns during monopolar medial stimulation across vertebral levels. Heatmaps show co-activation patterns at the lowest amplitude where loss of selectivity occurred during monopolar medial stimulation at vertebral levels L4, L5, L6, and L7. Rows represent nerves selectively recruited at threshold, and columns represent nerves that were subsequently co-activated and disrupted selectivity. The color of each cell reflects the ratio of co-activation events between each nerve pair. Red dashed lines separate sciatic and femoral nerve branches. Adjacent bar plots indicate how often each nerve was involved in recruitment loss as a disrupter nerve.

**Supplementary Figure 3:**
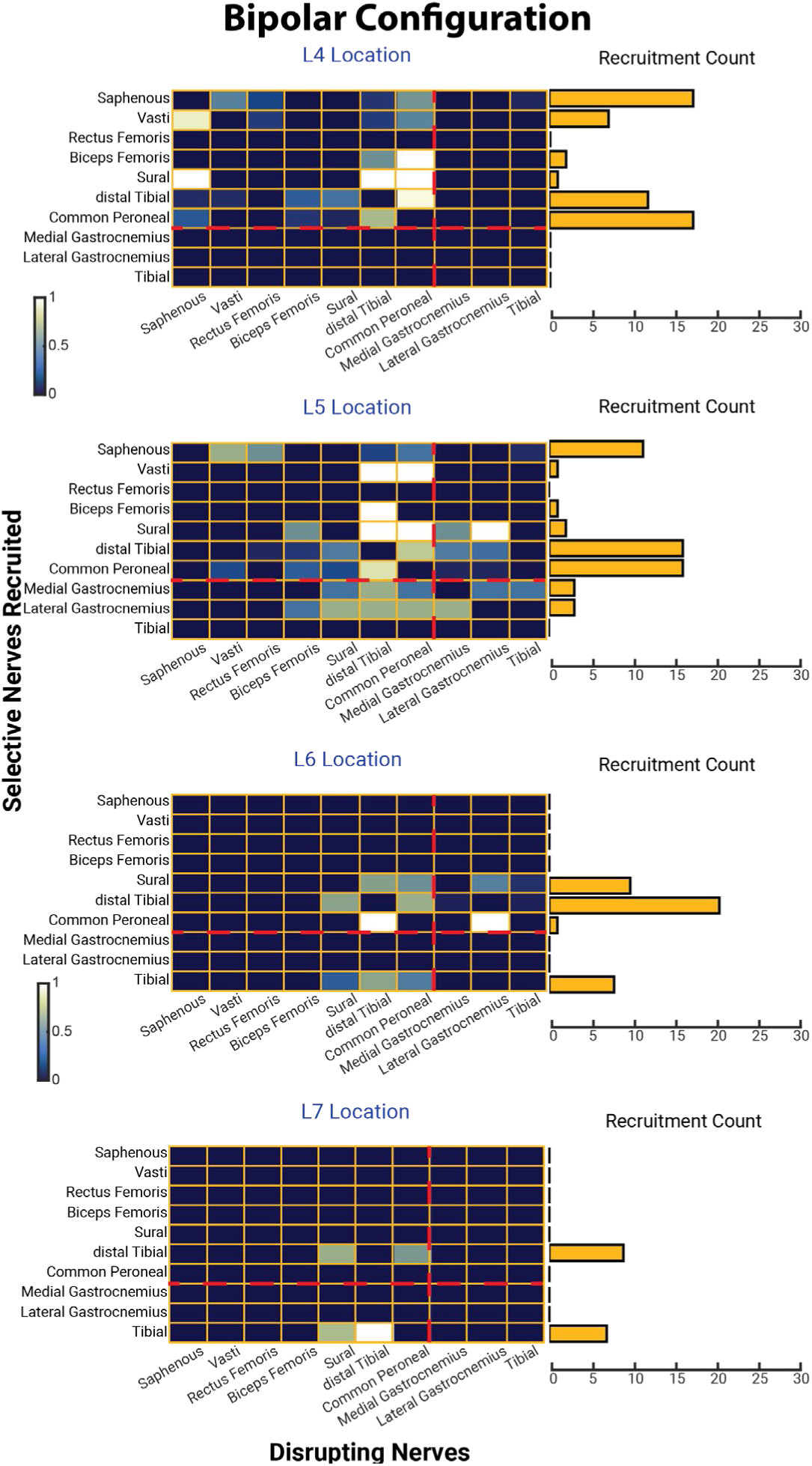
Co-activation patterns during bipolar stimulation across vertebral levels. Heatmaps showing loss of selectivity patterns during bipolar stimulation at vertebral levels L4, L5, L6, and L7. Rows represent nerves selectively recruited at threshold, and columns represent nerves that were subsequently co-activated and disrupted selectivity. The color of each cell reflects the ratio of co-activation events between each nerve pair. Red dashed lines separate sciatic and femoral nerve branches. Adjacent bar plots indicate how often each nerve was involved in recruitment loss as a disrupter nerve.

## Notes

### Competing Interest Statement

S.F.L. has equity in Hologram Consultants, LLC and is a member of the scientific advisory board for Abbott Neuromodulation. S.F.L. holds stock options, received research support, and serves on the scientific advisory board of Presidio Medical.

### Summary of Updates

We have added a citation (#23) to a companion manuscript that was also uploaded to bioRxiv.

### Funding

- National Institutes of Health: R01NS121028

